# Mitigating the Field-of-View – Resolution Tradeoff by Photon Superlocalization

**DOI:** 10.64898/2025.12.08.693070

**Authors:** Sulaimon Balogun, Andreas E. Vasdekis

## Abstract

Optical imaging systems are fundamentally constrained by a tradeoff between field of view (FoV) and spatial resolution. Long working-distance objectives, routinely used in biological imaging and especially in light-sheet microscopy, provide large FoV but reduced numerical aperture (NA) and magnification, broadening the point spread function (PSF) while coarsening detector sampling. As a result, even PSF-limited resolution is often undersampled. Here we demonstrate photon superlocalization as a strategy to mitigate this tradeoff. On intensified detectors, individual photons form multipixel detection clouds that can be centroided and reassigned to a finer virtual grid, thereby increasing the effective sampling frequency without optical modifications or FoV penalty. Proof-of-principle epifluorescence and light-sheet experiments show that photon superlocalization restores near-PSF-limited resolution and reveals subcellular structure otherwise obscured by undersampling. This approach provides a generalizable, photon-efficient pathway for improving spatial resolution across imaging modalities constrained by the FoV-resolution tradeoff.

## Introduction

Optical microscopes are fundamentally constrained by the Abbe diffraction limit, which defines the smallest resolvable distance between two adjacent features as approximately half the wavelength of light.^1, 2^ This limit, determined by the wave nature of light and the numerical aperture (NA) of the imaging objective, sets the ultimate bound of spatial resolution. In practice, however, most imaging systems operate below this limit due to sampling constraints that impose an innate trade-off between field of view (FoV) and resolution.^3^ Low-magnification, low-NA objectives, or equivalently detectors with large pixel sizes, provide a wide FoV but produce a broad point spread function (PSF) that is easily undersampled. Concomitantly, high-magnification, high-NA optics achieve fine resolution at the cost, however of FoV, photon efficiency (i.e., photons per pixel), and working distance.

This trade-off between FoV and spatial resolution represents a persistent challenge in modern microscopy of biological systems, where large-scale imaging and the resulting increased throughput, as well as minimal phototoxicity must be balanced against the need for subcellular detail.^4, 5^ This challenge is particularly evident in light-sheet fluorescence microscopy (LSM), also known as selective plane imaging microscopy (SPIM).^6–9^ In this context, long working distances, and inevitably low-NA detection optics, are needed to accommodate the stringent geometric requirement of two proximal objectives in related configurations. Achieving high-resolution, wide-area imaging without sacrificing photon efficiency remains a central objective in optical system design.

According to the Nyquist-Shannon sampling criterion, faithful image reconstruction requires at least two detector pixels per PSF full width at half maximum (FWHM).^10^ **Fig. 1a** illustrates this relationship by displaying the two-dimensional modulation transfer function (MTF) of an incoherent imaging system (λ = 600 nm, NA = 1.1, M = 40×), where the optical cutoff frequency (red circle) and the Nyquist limits for 6.5 µm (*green circle*, e.g., standard sCMOS detector) and 12.5 µm (*yellow circle*, e.g., standard EMCCD detector) pixel detectors delineate the sampled spatial band-width. As pixel size increases, the sampling frequency falls considerably lower than the optical cutoff, causing loss of high-frequency information. This effect is also visible in the radial MTF profile shown in **Fig. 1b**, which highlights the fidelity erosion when the sampling frequency approaches the Nyquist limit.

**Figure 1.**
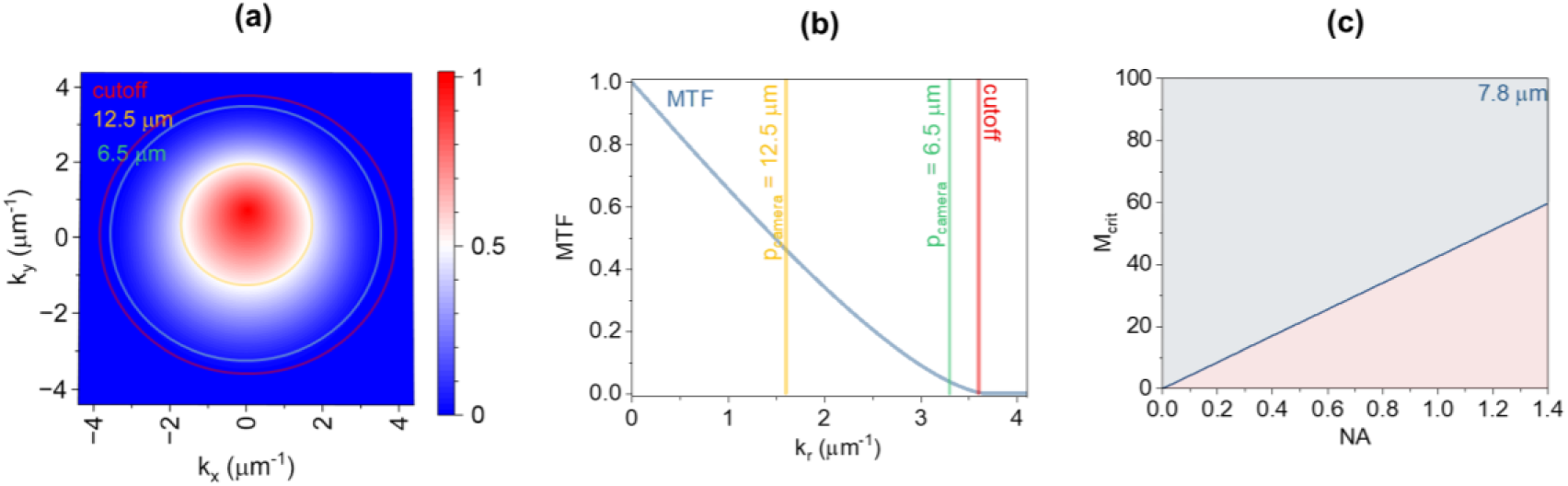
Modulation transfer and sampling limits. **(a)** Modulation transfer function (MTF) of an incoherent imaging system (λ = 600 nm, NA = 1.1, M = 40×). The diffraction-limited optical cutoff is shown in red, along with Nyquist frequencies for 6.5 µm (green) and 12.5 µm (yellow) detector pixels. **(b)** Radial MTF profile corresponding to the system parameters in **(a)**. **(c)** Nyquist sampling map as a function of objective magnification and numerical aperture for a 7.8 µm pixel detector, indicating undersampled (red) and oversampled (blue) operating regimes.

In practical imaging conditions, the effective sampling frequency of 5-7 µm pixels at 20×-40× magnification is often considerably lower than the optical cutoff, producing aliasing and loss of high-frequency detail.^11^ Increasing magnification or employing smaller-pixel sensors can alleviate undersampling, but only at the expense of FoV and signal-to-noise ratio, respectively. These constraints are particularly detrimental in bioimaging, where photon-limited conditions are key against photo-toxicity and photobleaching.^12–14^ The boundaries of this undersampling regime are mapped in **Fig. 1c**, which shows the Nyquist sampling cutoff as a function of objective magnification and NA for a 7.8 µm pixel detector (6.5 μm and 12.5 μm pixel sizes are shown in **Fig. S1**), with the undersampled and oversampled configurations separated by color (*red*-*blue*, respectively).

To address these limitations, we introduce *photon superlocalization*, a computational approach that effectively increases sampling density without modifying the optical hardware configuration. Instead of relying on the physical detector-pixel grid to define resolution, this method localizes individual photon detection events with subpixel precision, reconstructing an effective sampling lattice denser than the detector pitch.^15^ By exploiting the statistical distribution of photon arrivals, photon superlocalization recovers diffraction-limited spatial detail while preserving FoV and photon efficiency.

We have previously demonstrated a proof-of-concept implementation of this approach for Raman-scattered photons, while related methods have been reported in x-ray imaging, where they were described as superesolution.^16, 17^ Similar ideas were explored by applying single-molecule localization algorithms to improve photon centroiding accuracy.^18^ Here, we expand this framework to photon superlocalization in the broadly applicable domain of fluorescence imaging, including epi-and light-sheet fluorescence microscopy. In this context, we describe photon superlocalization as the recovery of photon positions beyond the detector sampling limit through statistical inference. Rather than a form of superesolution, our approach serves as a computational mitigation strategy in imaging that is sampling-limited. Through the analyses and experiments that follow, we demonstrate that photon superlocalization simultaneously enables high-resolution, wide-FoV imaging with conventional objectives, establishing a unified statistical framework across imaging modalities.

## Results

### Sampling Theory and Photon Superlocalization

Most optical imaging systems are constrained by the interplay between diffraction, sampling, and photon statistics. The lateral resolution (inversely related to the spatial frequency cutoff) of an incoherent, diffraction-limited imaging system is given by the Rayleigh criterion:^19^

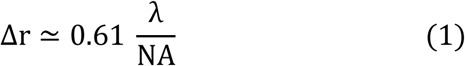

This relationship defines the smallest resolvable feature, which corresponds to the largest spatial frequency transmitted by the system’s MTF. However, digital imaging is additionally bound by the discrete sampling imposed by the detector. According to the Nyquist-Shannon criterion, the sampling period at the object plane, p_sample_ = p_camera_/M, must satisfy p_sample_ ≤ Δr/2 to avoid aliasing.^19^ Optical magnification M, thus, serves as the coupling term between the physical optics and the sampling architecture: increasing M improves sampling density but reduces the FoV, a persistent trade-off in wide-field microscopy.

The mutual dependence of optical cutoff, sampling frequency, and magnification is captured in **Fig. 1**. The two-dimensional MTF for an incoherent fluorescence imaging system (λ = 600 nm, M = 40 ×, NA = 1.1) defines the optical support bounded by the cutoff frequency f_c_ = 2·NA/λ. Superimposed Nyquist circles corresponding to 6.5 µm and 12.5 µm detector pixel sizes at 40× magnification illustrate how finite sampling constrains the transmitted spatial frequency bandwidth. As the sampling frequency approaches f_c_, the radial MTF (**Fig. 1b**) reveals the rapid collapse of transfer fidelity, noting that aliasing emerges well before the nominal cutoff is reached. Generalization across NA and magnification yields the critical magnification:

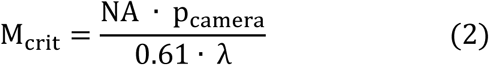

which separates undersampled regimes from Nyquist-satisfied ones (**Fig. 1c**). Collectively, these representations show that sampling losses are a coupled property of the optical-digital system: variations in NA or magnification redistribute the accessible spatial-frequency volume in tandem with the electronic sampling grid.

Photon superlocalization offers a physical way to circumvent this constraint by converting stochastic photon detection into an additional sampling channel. In intensified detectors, each absorbed photon triggers a localized avalanche whose spatial extent is typically asymmetric and covers several adjacent pixels, forming a cloud. The resulting cloud encodes the photon’s impact position with subpixel information content. Estimating the centroid of the photon-cloud yields a localization precision that, in the low-noise limit, approaches the Cramér-Rao bound, scaling approximately as:^20, 21^

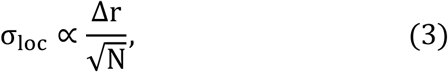

where Δr is the diffraction-limited width of the optical point-spread function and N is the number of detected photons comprising the cloud. When accumulated across many photon-clouds, these localized centroids effectively resample the image at a density far exceeding the native pixel grid, generating a more finely sampled representation of the diffraction-limited optical field.

### Approach

In this work, we employed an intensified CMOS (iCMOS) detector that inherently supports the processes mentioned in the previous subsection (**Methods**). Photons incident on the photocathode eject electrons whose trajectories depend both on impact position and incidence angle, producing slightly anisotropic emission footprints on the phosphor.^22, 23^ Within the microchannel plate (MCP), electron multiplication proceeds symmetrically about the trajectory axis, preserving a well-defined centroid even when the envelope of the charge cloud is elongated or skewed. The phosphor output, relayed through a fiberoptic taper to the CMOS sensor, therefore forms asymmetric light clusters whose centroids can remain statistically unbiased. This intrinsic asymmetry, illustrated in **Fig. 2**, allows each photon to be localized with precision below the detector’s pixel pitch, effectively transforming the intensifier into a hybrid over-sampler.

**Figure 2.**
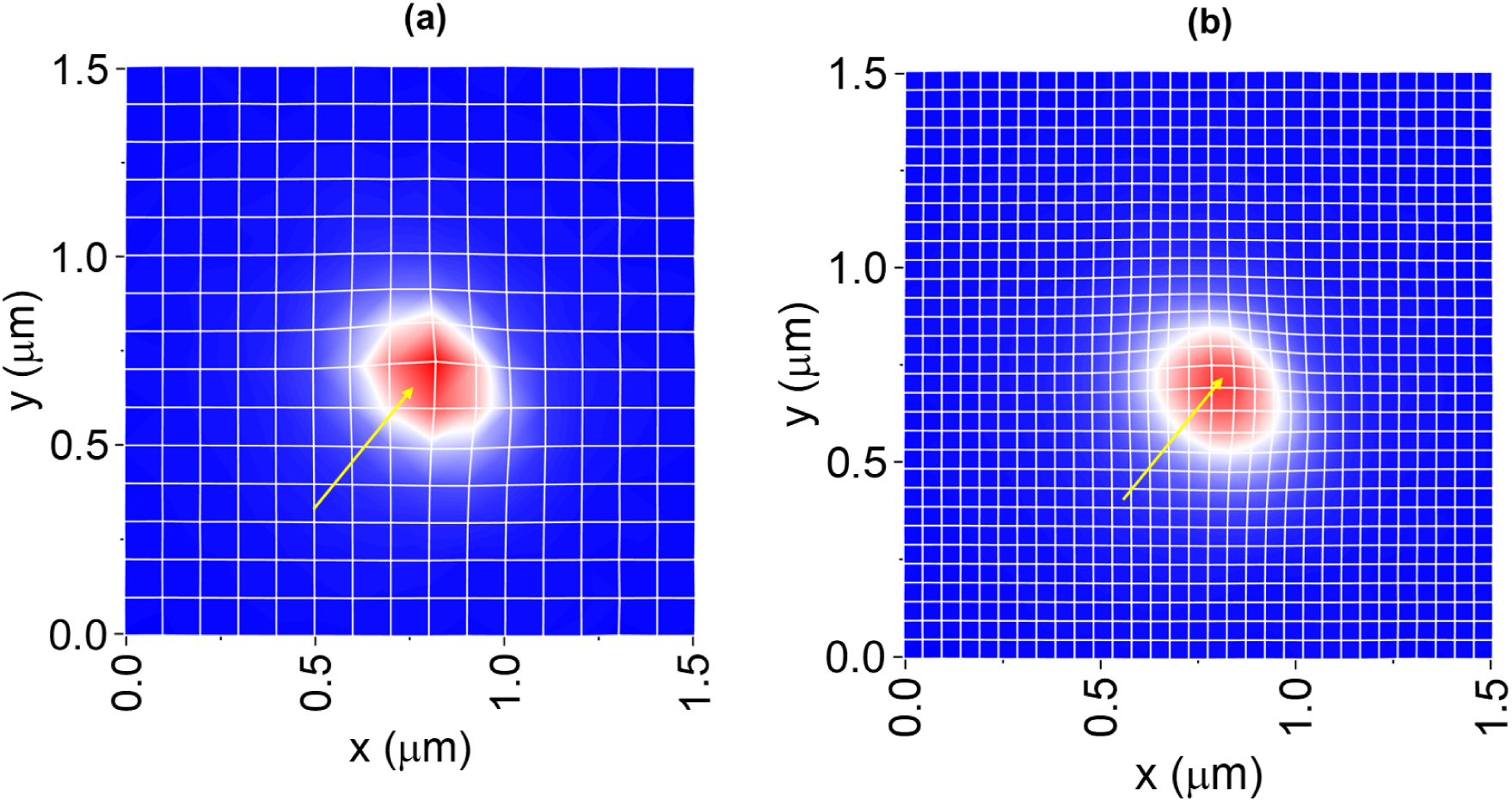
Photon superlocalization. **(a)** Native 1×1 detector sampling, where photon clouds (in *red*) and cloud centroids (*yellow* arrow) are assigned to the physical pixel grid. **(b)** 2×2 photon-superlocalized resampling in which photons are redistributed to a finer virtual grid to improve spatial sampling.

We acquired all images with the iCMOS detector operating at 100 kHz, while the detector’s gating electronics drove an electro-optic modulator (EOM) for illumination gating (**Methods**). The excitation wavelength was 561 nm, delivered through a standard inverted microscope (Leica DMi8). For epifluorescence imaging, we coupled the excitation beam through the detection objective, whereas for LSM/SPIM we employed the same microscope platform with a modified sample-stage to enable selective-plane illumination without altering the detection path, as we have reported previously.^16^ The EOM temporally synchronized the epi- or light-sheet excitation with frame acquisition, ensuring one-to-one correspondence between photon arrival statistics and detector readout. In addition to cooling the sensor down to -25°C, this synchronization suppressed dark counts to approximately 6·10^-7^ ± 8·10^-8^ (mean ± SEM), enabling reliable photon-cloud segmentation at 3.6 kHz CMOS frame rates.

We extracted photon-cloud centroids using a maximum-entropy segmentation and weighted-moment analysis pipeline. Each frame was first thresholded using a global maximum-entropy criterion to isolate statistically significant photon clouds, following mild top-hat background correction to suppress slow-varying illumination gradients. The resulting binary masks were refined by two sequential morphological erosions and size-based filtering to remove spurious detections and merge fragmented photon events. For each surviving component, intensity-weighted centroids were computed directly from the raw pixel intensities, producing per-frame centroid maps at progressively finer virtual grids (1× and 2× relative to the native pixel size of 7.8 μm). Each centroid map subsequently underwent denoising using the PURE-LET algorithm,^24^ followed by median filtering to suppress isolated outliers and Gaussian blurring to enforce continuity of the reconstructed intensity field. This processing sequence preserved the statistical structure of the photon-distribution while enhancing spatial coherence, yielding superlocalized representations suitable for subsequent resolution and convergence analyses. Finer sampling factors (3× and 4×) suffered from photon-number limitations and the diminishing returns due to oversampling, as further detailed below.

This weighted-centroid method implements a discrete form of the mass-centroid estimator, where pixel intensities serve as statistical weights proportional to local photon density. Under Poisson noise, the centroid-mediated mean intensity follows the expected theoretical scaling with temporal variance, as detailed below for each imaging condition. Summing thousands of such localized photon-clouds produced images whose effective sampling frequency approached and eventually exceeded the Nyquist threshold shown in **Fig. 1c**. Importantly, this occurred without any modifications to the optical path or reductions in the imaging FoV. In doing so, the iCMOS detector effectively re-parameterized the sampling process, enhancing the imaging spatial information content and bridging the gap between optical and digital resolution through photon superlocalization.

### Imaging Assessment

#### Particles

We first implemented photon superlocalization on a conventional epifluorescence microscope equipped with a 40×/1.1 NA objective and the iCMOS detector (7.8 μm pixel size). Under these conditions, the system operated below the Nyquist sampling requirement for the diffraction-limited point-spread function (PSF), resulting in undersampling of fine spatial frequencies (**Fig. 1c**). To assess localization performance across spatial regimes, we imaged fluorescent particles of 100 nm and 1 µm diameters and reconstructed their intensity profiles to compute the full width at half maximum (FWHM). For the 100 nm particles, the FWHM decreased from approximately 1.46 µm ± 0.05 μm at native sampling (1×1, mean ± SEM) to 1.15 µm ± 0.05 μm after 2×2 photon-superlocalized resampling (**Fig. 3a**), indicating an improvement in spatial resolution towards the resolution limit defined by the optical PSF. We observed no further resolution improvements at 3×3 and 4×4 finer grids, conditions that lead to oversampling at the cost of the average number of photons per pixel (**Fig. S2**).

**Figure 3.**
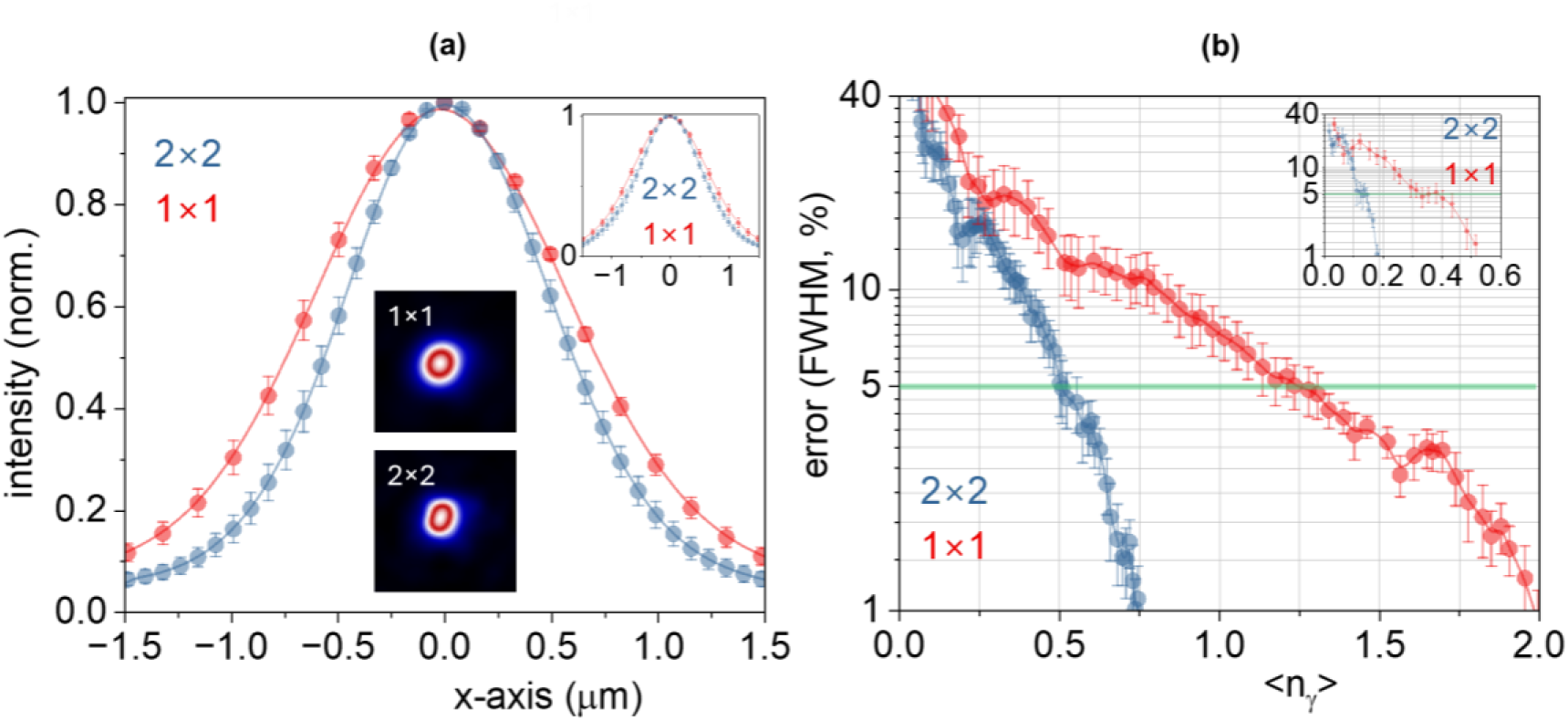
Resolution improvement with photon superlocalization. **(a)** Line profiles of 100 nm particles showing enhanced apparent resolution using 2×2 resampling (blue) compared with native 1×1 sampling (red) (for *n = 8* particles, points and error bars represent the mean ± SEM). *Insets:* 1 µm particle intensity profile (*top right*) and representative 100 nm particle images for both sampling modes (*bottom*). **(b)** Localization precision as a function of photons per pixel, demonstrating faster convergence for 2×2 reconstruction (for *n = 8* particles). Green line denotes the 5% error threshold, while *inset* shows convergence behavior for 1 µm particles.

We observed a comparable resolution improvement for the 1 µm particles (1.58 µm ± 0.08 µm at native 1×1 sampling to 1.29 µm ± 0.08 µm after 2×2 photon-superlocalized resampling; **Fig. 3a**, *inset*). For both particle sizes, the 2×2 reconstructions exhibited faster convergence of localization precision with photon-per-pixel count (**Fig. 3b**). This earlier convergence is not due to an increased number of detected photons but rather reflects a redistribution of the same photons across a finer virtual sampling grid. Photon-cloud centroids provide subpixel positional information, allowing each photon to contribute to a more densely sampled representation of the emitter. As a result, the effective sampling density increases even though the total photon budget remains unchanged.

#### Bioimaging

We next extended the approach to biological imaging using two representative systems: fixed plant roots and fixed yeast cells, to evaluate performance across differing structural and dynamic regimes. For *Medicago truncatula* roots, fixed and stained with propidium iodide (PI), epifluorescence imaging was performed using a 4×/0.10 NA objective and the iCMOS detector. This optical configuration lies well below Nyquist sampling, resulting in coarse representation of root boundaries and intracellular fluorescence intensity (**Fig. 1c**). Finer sampling yielded a more accurate representation of the same features and enabled earlier convergence with respect to photon flux (**Fig. S3**). To quantify this improvement, we employed image decorrelation analysis.^25^ Two-dimensional Fourier transforms of the root images were radially averaged, revealing a clear resolution improvement in the 2×2 condition, manifested as an increased (decorrelation) cutoff frequency (k_c, 1×1_ = 0.13 ± 0.003 μm^-1^ and k_c, 2×2_ = 0.29 ± 0.004 μm^-1^, **Fig. 4a**, **b** and **Fig. S4**).

**Figure 4.**
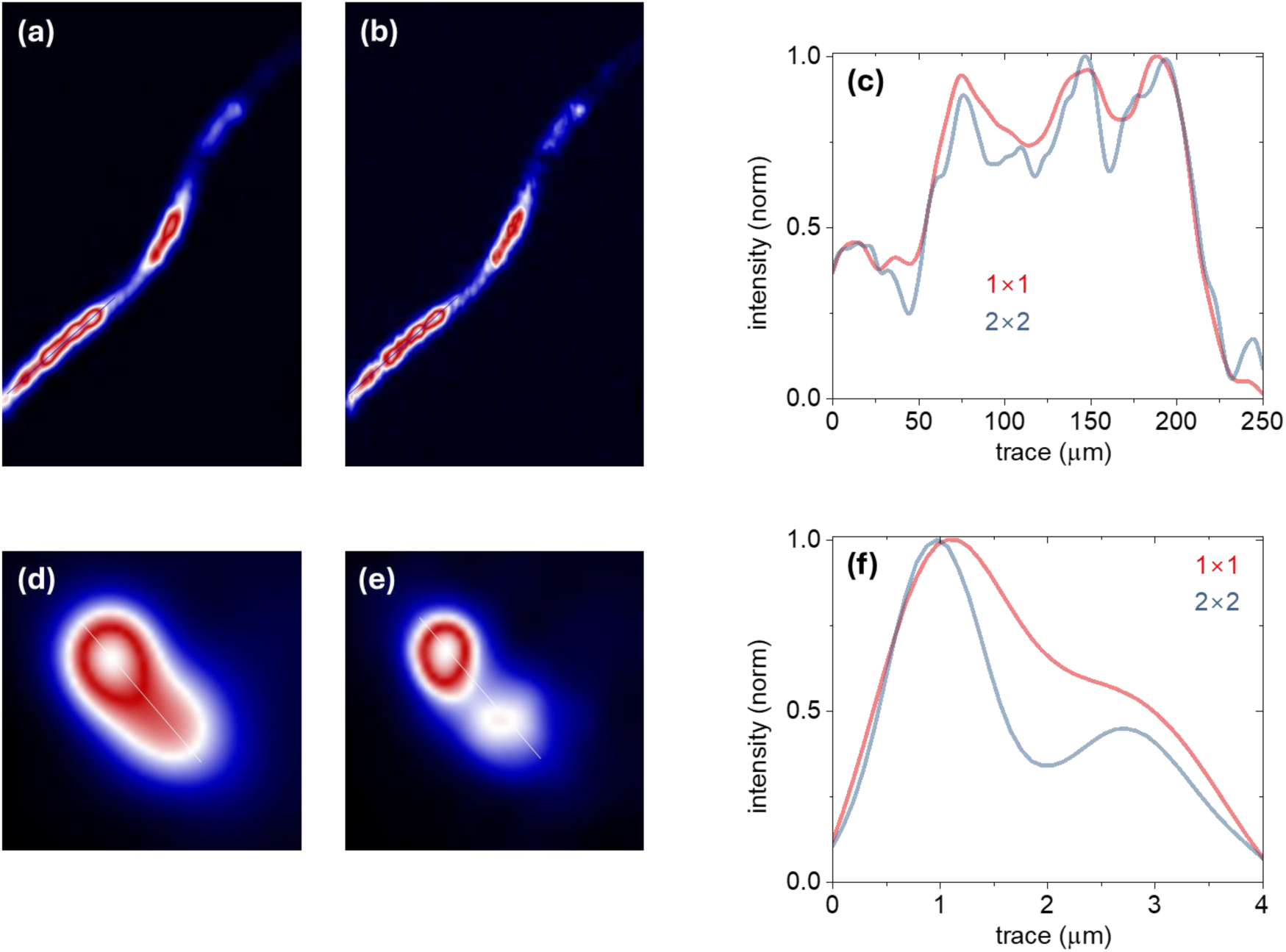
Enhanced resolution in biological samples using 2×2 resampling. **(a)** Native 1×1 sampling of plant root structures. **(b)** 2×2 reconstructed image showing improved visibility of fine root features while maintaining the original field of view. **(c)** One-dimensional intensity trace along the indicated region. **(d)** Native 1×1 sampling and **(e)** 2×2 resampling that enhances cytosolic contrast, with the corresponding one-dimensional intensity trace **(f)**.

The improvement demonstrates that photon superlocalization effectively compensates for optical undersampling in extended biological specimens. Qualitatively, applying photon superlocalization with 2×2 resampling (**Fig. 4b** and **Fig. S4**) enhanced the delineation of fine root structures while preserving the native FoV observed under conventional 1×1 sampling. Although subcellular boundaries were not individually resolved, the 2×2 reconstruction revealed greater structural continuity and sharper contrast along elongated regions. Insets in Fig. **4a**, **b** (and **Fig. S4**) show representative intensity profiles highlighting increased feature content and finer spatial modulation in the 2×2 dataset.

For single-cell light-sheet fluorescence microscopy (LSM) of *Yarrowia lipolytica* stained with Nile-Red, we applied photon superlocalization to datasets acquired using a 40×/1.1 NA illumination objective detection objective and a 20×/0.42 NA illumination objective (**Methods**). In this configuration, the native 1×1 intensity images primarily reflected global illumination gradients across the cell (**Fig. 4d** and **Fig. S5**), whereas the 2×2 photon-superlocalized reconstructions (**Fig. 4e** and **Fig. S5**) revealed sharper axial transitions with enhanced contrast on the cell cytosol. Insets in **Fig. 4c** and **d** display representative intensity traces, showing that the 2×2 dataset exhibits additional local intensity modulations consistent with improved spatial frequency recovery. Quantitative decorrelation analysis confirmed these observations: the 2×2 reconstructions exhibited a reproducible increase in k_c_ (Δk_c_ = 0.05 ± 0.01) relative to the native 1×1 data, corresponding to a considerable enhancement in resolvable spatial frequency.

To further assess the noise regime, we performed mean-variance analyses of pixel intensities across repeated acquisitions. The resulting linear relationship between mean signal and variance (**Fig. 5**) confirmed that the system operated at the Poisson, or shot-noise, limit. This validation establishes that the observed k_c_ gains reflect improvements in photon sampling efficiency rather than hardware or illumination artifacts. As with the particle and root datasets, the earlier k_c_ convergence in the 2×2 condition (**Fig. S3**) thus arises from more efficient reallocation of photon statistics within the same total photon budget, emphasizing that the benefit derives from optimized photon utilization rather than increased excitation or emission yield.

**Figure 5.**
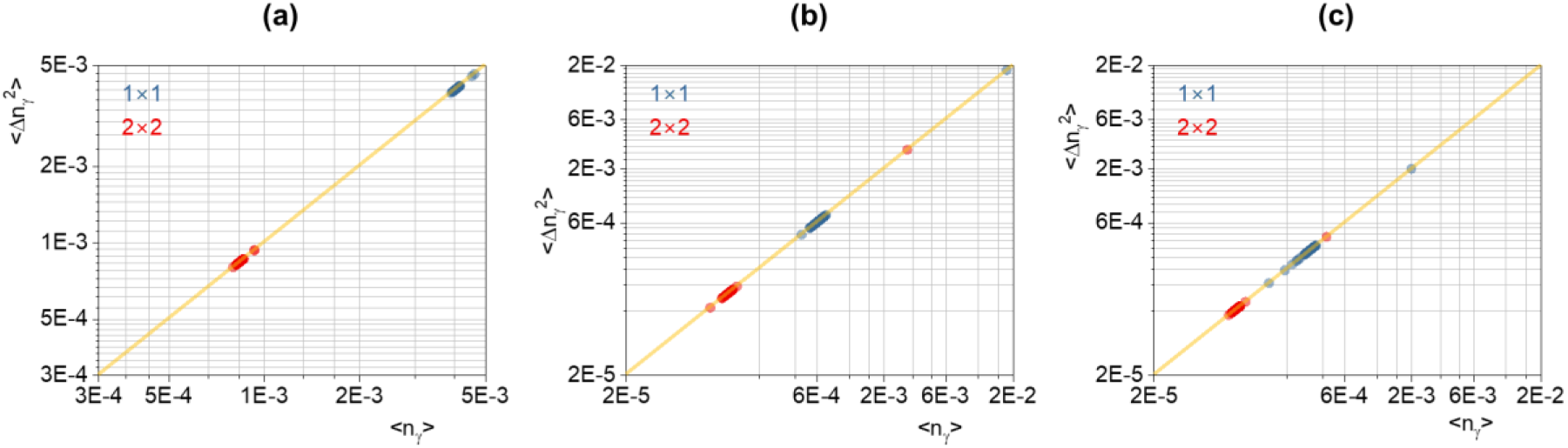
Shot-noise-limited photon statistics across samples. Mean–variance analysis for yeast **(a)**, 100 nm diameter particles **(b)**, and **(c)** plant root samples, showing a linear mean-variance relationship consistent with shot-noise-limited detection.

## Discussion

Widefield and light-sheet fluorescence imaging often operate under sampling-limited conditions where moderate-NA, long-working-distance objectives broaden the PSF beyond what detector pixel sizes can adequately sample. This mismatch leaves a substantial portion of the optical bandwidth unrecorded, even when photon budgets would otherwise permit higher resolution. Photon superlocalization provides a way to bridge this gap by converting the intrinsic structure of photon-cloud footprints into additional sampling information, effectively increasing the density of measurements without altering the optical path, FoV, or photon load.

Across all tested systems, ranging from subwavelength particles, yeast cells, and plant roots, we found that 2×2 resampling consistently recovered higher spatial frequencies and improved structural fidelity relative to native sampling. These gains manifested as narrower particle profiles, increased decorrelation-based cut-off frequencies, and earlier convergence of resolution metrics under identical photon budgets. The improvement arises not from additional photons but from more efficient allocation of existing photons into finer virtual sampling bins, leveraging the localization precision permitted by Poisson arrival statistics.

Importantly, photon superlocalization does not extend beyond the diffraction-limited bandwidth. This explains the diminishing returns of 3×3 and 4×4 grids: once the virtual grid sufficiently samples the optical transfer function, further refinement probes unsupported spatial frequencies and cannot yield additional information. Thus, the method should be understood as a means to restore diffraction-limited performance in undersampled systems, not as a route to super-resolution.

Overall, photon superlocalization provides a broadly applicable, photon-efficient strategy for retrieving spatial information lost to coarse detector sampling. It offers particular value for geometrically constrained modalities such as LSM/SPIM, where increasing NA or magnification is often infeasible. By repurposing the physics of intensified detection, this approach expands the practical resolution of large-FoV fluorescence imaging without imposing hardware tradeoffs, illumination penalties, or complexity in optical design.

## Acknowledgments

We gratefully acknowledge funding from the U.S. Department of Energy, Office of Science, Office of Biological and Environmental Research (DE-SC0025418). We also acknowledge Maria J. Harrison (Boyce Thompson Institute) for preparing and providing the *Medicago truncatula* samples used in this research.

## Contributions

S.B. performed experiments and data analyses. A.E.V. performed centroid computations, supervised the project, and drafted the manuscript.

## Data Availability

The datasets generated and analyzed during this study are available from the corresponding author upon reasonable request. The MATLAB scripts used in this study are currently hosted in a private GitHub repository (https://github.com/aevasdekis/photon-superlocalization) and will be made publicly accessible upon publication.

## Methods

### Samples

To evaluate and validate the performance of photon superlocalization, we examined three distinct sample types: fluorescent particles, yeast cells, and plant roots.

#### Fluorescent particles

We imaged fluorescent polystyrene beads of 100 nm and 1000 nm diameter (Bangs Laboratories), which served as point-like emitters for assessing spatial resolution and localization behavior.

#### Yeast cells

The second sample comprised *Yarrowia lipolytica* (strain FEB-168). Cells were revived from a frozen glycerol stock (-80 °C) onto YPD plates and subsequently cultured twice in YPD for 24 h at 28 °C (20 mL cultures in 125 mL shaker flasks). A second passage was transferred into defined medium (YSM) and grown for 66 h under the same shaking conditions. Following culture, cells were fixed overnight in 2% paraformaldehyde, washed three times in PBS, and stained with Nile Red (1 µg/mL in DMSO; Thermo Fisher). YPD medium consisted of 20 g/L Bacto Peptone (BD), 10 g/L yeast extract (Alfa Aesar), and 20 g/L glucose (Fisher). YSM medium was prepared using 1.7 g/L yeast nitrogen base without amino acids and ammonium sulfate (BD Difco), 0.69 g/L CSM without leucine (Sunrise Science Products), 0.1 g/L leucine, 1.1 g/L ammonium sulfate (Fisher), and 50 g/L glucose (Fisher).

#### Plant roots

The third sample consisted of *Medicago truncatula* (R108) roots. Plants were grown for 4 weeks in a sterile sand/gravel substrate under a 16 h light (25 °C) / 8 h dark (22 °C) cycle and fertilized twice weekly with modified half-strength Hoagland’s solution.^26^ Root tissues were fixed in 50% ethanol (v/v), and root tips and secondary branches were excised and stained with propidium iodide (1 µg/mL; Invitrogen).

### Experiment setup for particles and roots imaging

For particle and root imaging, we employed an epifluorescence microscope. Illumination was provided by a continuous-wave 561 nm diode laser (OBIS, Coherent; 200 mW average power), whose output was stabilized using a motorized attenuator in a closed-loop configuration (VA-BB-2-CONEX, Newport). The beam was modulated at 100 kHz using an electro-optic modulator (LM0202, Excelitas) that was driven directly by the gating electronics of the intensified CMOS (iCMOS) detector. Illumination was delivered in a collimated format by combining a 180 mm lens with either a 4×/0.1 NA objective (HI PLAN, Leica) for *M. truncatula* roots or a 40×/1.1 NA objective (HC PL APO, Leica) for fluorescent particles, forming a 4f configuration that produced a Gaussian excitation profile. Under these conditions, specimens received ∼5 pW/µm^2^ irradiance, enabling ≤1 photon-per-pixel events in a multishot acquisition regime. Fluorescence detection was performed using a 40×/1.1 NA objective that relayed the emission onto an intensified CMOS camera (HiCAM Fluo, Lambert Instruments). The iCMOS comprises a 1280 × 1024 CMOS sensor (7.8 µm pixel pitch), a double-stage microchannel plate (MCP), and a GaAs photocathode with P46 phosphor, providing an overall quantum efficiency (QE) of ∼30%. The detector inter-faced with a workstation (128 GB RAM) via a four-channel CoaXPress connection. To benchmark performance, identical specimen regions were simultaneously projected onto both the iCMOS and a scientific CMOS camera (sCMOS, ORCA-Flash 4.0 v2, Hamamatsu) under the same illumination. While the sCMOS provided ∼82% QE, the ∼30% QE of the iCMOS highlights the potential for further gains in near-zero-photon imaging as intensified-sensor architectures continue to advance. Illumination was synchronized to the intensifier to minimize background light: the 100 kHz modulation was triggered by the intensifier gate, resulting in ∼25 intensifier firings per CMOS frame. To immobilize samples and maintain hydration during imaging, we prepared a low-temperature agarose substrate. Agarose (1.2% w/v; Fisher Scientific) was dissolved in the appropriate medium, deionized water for *M. truncatula* roots and 1× PBS for fixed *Y. lipolytica* cells and fluorescent particles, by heating to 80 °C for 45 min. After cooling to ∼35 °C, the molten agarose was cast between two coverslips and allowed to solidify for ∼15 min, forming a uniform ∼500 µm-thick pad. Particles and biological samples were placed on the agarose surface and covered with a clean coverslip to ensure stable and reproducible optical conditions.

### Experimental setup for yeast cell imaging

We used a fluorescence light-sheet imaging (LSM) configuration integrated into a standard inverted microscope, as we recently reported.^16^ The microscope was controlled through Micro-Manager,^27^ with excitation delivered through a 20×/0.42 NA objective to generate a ∼2 µm diameter Gaussian beam.^16^ A MEMS mirror laterally scanned the beam to form images with fields of view up to 200 µm × 10 µm. Continuous-wave 561 nm excitation was modulated at 1 kHz using a rotating disk chopper and synchronized to the intensified CMOS detector (iCMOS) to suppress background illumination. Under these conditions, specimens received irradiation levels as low as 20 nW/μm^2^, enabling <1 photon-per-pixel events in a multishot acquisition format. The iCMOS detector operated with 5 µs gate widths, 100 kHz gating rates, and 1 kHz CMOS readout rates. At these settings, the intensifier fired 50 times per CMOS frame. Single-photon images were stored in 12-bit format in system RAM, and subsequent processing was performed on 16-bit TIFF files saved to disk.

### Photon Cloud Centroid Estimation

Photon-cloud centroids were extracted using a maximum-entropy segmentation and weighted-moment pipeline in MATLAB. Raw iCMOS frames were optionally background corrected and mildly opened before applying a global maximum-entropy threshold to identify photon-cloud candidates. The resulting binary mask underwent two erosions and minimum-area filtering, after which connected components were screened using geometric (area, solidity, eccentricity, circularity) and intensity criteria. Components passing these vetoes were treated as valid photon clouds. Weighted centroids were then computed using the raw intensities and projected onto virtual sampling grids (1×1, 2×2, 3×3, 4×4) by scaling centroid coordinates to the corresponding grid size. Each frame thus yielded centroid maps at progressively finer effective sampling densities.

### Data analysis

#### Full-Width-at-Half-Maximum (FWHM)

FWHM were computed by fitting the trace of each particle with a Gaussian function using Origin Pro.

#### Statistics

All other statistics, such as mean, and SEM were computed in Origin Pro.

#### Errors – Convergence

For particle data, we acquired image sequences in which each dataset comprised 6000 frames. The final frame in each sequence was treated as the ground-truth reference. FWHM values were computed for each frame prior to calculating the percent error relative to the ground truth. For biological samples (*Y. lipolytica* cells and *M. truncatula* roots), we quantified resolution using decorrelation analysis.^25^ Two-dimensional Fourier transforms of each image were radially averaged to obtain the decorrelation-based cutoff frequency *k*_*c*_, after which the percent error was computed analogously.

The percent error for either metric (*k*_*c*_ or FWHM) was defined as:

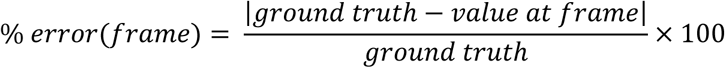

## References

1. Abbe, E. Beiträge zur Theorie des Mikroskops und der mikroskopischen Wahrnehmung. Archiv für Mikroskopische Anatomie 9, 413–468 (1873).

2. Hell, S.W. Far-field optical nanoscopy. Science (New York, N.Y.) 316, 1153–1158 (2007).

3. Montgomery, W.D. Sampling in imaging systems. J. Opt. Soc. Am. 65, 700–706 (1975).

4. Yi, C. et al. Video-rate 3D imaging of living cells using Fourier view-channel-depth light field microscopy. Communications biology 6, 1259 (2023).

5. Wu, Y. & Shroff, H. Multiscale fluorescence imaging of living samples. Histochemistry and cell biology 158, 301–323 (2022).

6. Vettenburg, T. et al. Light-sheet microscopy using an Airy beam. Nature methods 11, 541–544 (2014).

7. Stelzer, E.H.K. et al. Light sheet fluorescence microscopy. Nature Reviews Methods Primers 1, 73 (2021).

8. Chen, B.-C. et al. Lattice light-sheet microscopy: Imaging molecules to embryos at high spatiotemporal resolution. Science (New York, N.Y.) 346, 1257998 (2014).

9. Kasu, R., Luo, H., Luckhart, S., Christodoulides, D.N. & Vasdekis, A.E. Single-Objective Airy Light-Sheet Imaging. ACS Photonics 12, 2432–2439 (2025).

10. Goodman, J.W. Introduction to Fourier optics. (Roberts and Company publishers, 2005).

11. Dobbins, J.T., 3rd Effects of undersampling on the proper interpretation of modulation transfer function, noise power spectra, and noise equivalent quanta of digital imaging systems. Medical physics 22, 171–181 (1995).

12. Icha, J., Weber, M., Waters, J.C. & Norden, C. Phototoxicity in live fluorescence microscopy, and how to avoid it. BioEssays : news and reviews in molecular, cellular and developmental biology 39 (2017).

13. Young, I.T., Gerbrands, J.J. & Van Vliet, L.J. Fundamentals of image processing. (1998).

14. Hoebe, R.A. et al. Controlled light-exposure microscopy reduces photobleaching and phototoxicity in fluorescence live-cell imaging. Nature Biotechnology 25, 249–253 (2007).

15. Small, A. & Stahlheber, S. Fluorophore localization algorithms for super-resolution microscopy. Nature methods 11, 267–279 (2014).

16. Dunn, L. et al. Video-rate Raman-based metabolic imaging by Airy light-sheet illumination and photon-sparse detection. Proceedings of the National Academy of Sciences 120, e2210037120 (2023).

17. O’Connell, D.W. et al. Photon-counting, energy-resolving and super-resolution phase contrast X-ray imaging using an integrating detector. Opt. Express 28, 7080–7094 (2020).

18. Hirvonen, L.M., Barber, M.J. & Suhling, K. Photon counting imaging and centroiding with an electron-bombarded CCD using single molecule localisation software. Nuclear Instruments and Methods in Physics Research Section A: Accelerators, Spectrometers, Detectors and Associated Equipment 820, 121–125 (2016).

19. Born, M. & Wolf, E. Principles of optics: electromagnetic theory of propagation, interference and diffraction of light. (Elsevier, 2013).

20. Thompson, R.E., Larson, D.R. & Webb, W.W. Precise Nanometer Localization Analysis for Individual Fluorescent Probes. Biophysical Journal 82, 2775–2783 (2002).

21. Descloux, A.C., Grußmayer, K.S. & Radenovic, A. Parameter-free rendering of single-molecule localization microscopy data for parameter-free resolution estimation. Communications biology 4, 550 (2021).

22. Hadfield, R.H. Single-photon detectors for optical quantum information applications. Nature Photonics 3, 696–705 (2009).

23. Buller, G.S. & Collins, R.J. Single-photon generation and detection. Measurement Science and Technology 21, 012002 (2010).

24. Luisier, F., Vonesch, C., Blu, T. & Unser, M. Fast interscale wavelet denoising of Poisson-corrupted images. Signal Processing 90, 415–427 (2010).

25. Descloux, A., Grußmayer, K.S. & Radenovic, A. Parameter-free image resolution estimation based on decorrelation analysis. Nature methods 16, 918–924 (2019).

26. Floss, D.S., Levy, J.G., Lévesque-Tremblay, V., Pumplin, N. & Harrison, M.J. DELLA proteins regulate arbuscule formation in arbuscular mycorrhizal symbiosis. Proceedings of the National Academy of Sciences 110, E5025–E5034 (2013).

27. Edelstein, A.D. et al. Advanced methods of microscope control using μManager software. Journal of biological methods 1 (2014).

